# Generating, curating, and evaluating *trnL* reference sequence databases: Benchmarking OBITools3/ecoPCR, RESCRIPt, and MetaCurator

**DOI:** 10.64898/2026.04.07.717010

**Authors:** Ozge S. Kuddar, Kelly A. Meiklejohn, Benjamin J. Callahan

## Abstract

Plant DNA metabarcoding enables the identification of plant taxa in mixed samples, with the *trnL* (UAA) intron and its P6 loop mini-barcode region performing as well as or better than other commonly used markers. Reliable metabarcoding requires high-quality reference databases, yet a regularly maintained *trnL* resource is currently lacking. Consequently, most studies use uncurated sequences downloaded directly from public repositories without essential validation. We address these gaps by providing guidance through a systematic comparison of three database curation tools - OBITools3/ecoPCR, RESCRIPt, and MetaCurator - to generate three *trnL* reference sequence databases and evaluate their classification performance across commonly sequenced *trnL* regions (CD, CH, and GH).

Reference *trnL* sequences and taxonomy files were retrieved from public sequence repositories and curated using standardized filtering steps to reduce taxonomic errors, sequence ambiguity, and redundancy. Four simulated query datasets—two base sets and their mutated counterparts—were constructed to assess classification performance of the databases using the Naïve Bayesian Classifier implemented in DADA2.- The evaluation showed that performance differed by *trnL* region: MetaCurator and RESCRIPt yielded higher and similar metrics for *trnL* CD; OBITools3/ecoPCR and RESCRIPt were comparable for *trnL* CH; and MetaCurator attained the highest performance for *trnL* GH region. All reference databases, taxonomy, and evaluation files are available at Zenodo (https://doi.org/10.5281/zenodo.17969450). The complete computational workflow and scripts are available on GitHub (https://github.com/oskuddar/trnL_DB). Although evaluation was focused on plant taxa in the United States, the resulting databases are suitable for use as global *trnL* reference databases.

## 1. Introduction

Plant DNA metabarcoding – the amplification and sequencing of short, standardized DNA regions (barcodes) – is widely applied to analyze the abundance and diversity of plant taxa in mixed samples. Metabarcoding has been used to study large-scale plant diversity and community composition (Vasar et al., 2023), contemporary vegetation biodiversity in lake sediments (Alsos et al., 2018), compositional changes in vegetation across space and time in Arctic samples (Willerslev et al., 2014), plant–pollinator interactions (Bell et al., 2017), and dietary plant diversity in humans (Petrone et al., 2022) and animals (Yang et al., 2016). Metabarcoding has been applied to practical problems including food and plant-based medicine authentication (Arulandhu et al., 2019) and forensic investigations (Paranaiba et al., 2019).

Commonly used plant barcode regions include *matK*, *rbcL*, *trnH-psbA*, and *ycf1* (Dong et al., 2012) and *trnL* (Taberlet et al., 2007), all located in the chloroplast genome, as well as ITS in the nuclear genome. These regions vary in their interspecific (between species) and intraspecific (within species) resolution, affecting their ability to distinguish plant taxa. While multiple studies (CBOL Plant Working Group et al., 2009; Wizenberg et al., 2023; Richardson et al., 2015) have proposed the use of a multi-locus barcoding approach for achieving higher taxonomic resolution of plant communities, most metabarcoding studies rely on a single locus and selection remains dependent on the research objective, DNA sample quality, the coverage and representation of the database used to assign sequences, and the preferred sequencing method.

Commonly sourced sample types that are subjected to plant DNA metabarcoding include environmental matrices (e.g., soil, water, feces, dust air), archaeological and permafrost samples, and processed foods. Due to exposure to prolonged environmental conditions or chemical and mechanical processing, such samples often contain degraded DNA. Highly degraded (fragmented) DNA is nicked and depurinated, making it challenging to amplify and sequence barcoding regions, as the amplifiable fragment length progressively shortens with degradation level. Consequently, the use of mini-barcodes (typically <200 bp) has increased, since shorter regions are also more likely to be successfully amplified from degraded samples. However, obtaining accurate species-level classification with mini-barcodes represents a challenge, as shorter sequences can often contain limited genetic information. There is no agreement on the ideal mini barcode region to permit species level classification for most plant taxa, but the chloroplast *trnL* region has been suggested to perform better in terms of amplification efficiency and classification in specific applications. Thus, in this study, we focus on the chloroplast *trnL* (UAA) region, a Group I intron in the chloroplast genome, first introduced as a plant barcode by Taberlet et al. (1991) and further elaborated in (2007), characterized by its conserved secondary structure with alternating conserved and variable regions.

The *trnL* P6 loop outperformed *rbcL* mini-barcodes (rbcLZ1 and rbcL19b) in classifying plant communities in animal diet studies (Mallott et al., 2018), and it also performed better than ITS2 when applied to pollen from honey and soil eDNA (Milla et al., 2021; Ariza et al., 2024). Similarly, the *trnL* (UUA) intron was more effective than *rbcL* and *matK* for identifying xerothermic plants in Poland (Heise et al., 2015). *In situ* classification assessed using Naïve Bayesian Classifier identified *trnL* and *matK* as markers with genus-level resolution that outperformed *rpoB*, *rbcL*, *psbA-trnH*, and *psbK* (Matiz-Ceron et al., 2022).

The barcoding region is not the sole factor determining success in plant DNA metabarcoding studies; classification also depends on the quality, sequence diversity, and coverage of reference databases used for comparison of unknown sequences to permit taxonomic assignment. Despite the availability of both global (CALeDNA: Curd et al., 2019; MetaCurator: Richardson et al., 2020; rCRUX: Curd et al., 2024) and local (South African savanna: Botha et al., 2023; UHURU: Gill et al., 2019; Taohongling Sika Deer National Nature Reserve: Liu et al., 2024) *trnL* reference databases, maintaining them remains a significant challenge. As a result, many researchers rely on keyword searches to retrieve up-to-date sequences for specific genetic regions from publicly available databases such as GenBank (Sayers et al., 2025) and RefSeq (Goldfarb et al., 2025). However, sequences in databases like NCBI’s (National Center for Biotechnology Information) GenBank are user-submitted and issues often arise with metadata including typos, incomplete or incorrect annotations, or entirely missing information (Chorlton, 2024). Moreover, broad keyword searches may not retrieve genomic DNA sequences that contain the region of interest.

While sequence retrieval might be a relatively straight-forward process, curation of acquired sequences is essential to ensure reliable downstream taxonomic assignment. Key curation steps include assigning sequences to the most up-to-date taxonomic information (e.g., NCBI Taxonomy, which is updated hourly), dereplicating by taxon ID, removing sequences with ambiguous bases exceeding a user-defined threshold, filtering by expected amplicon length, and excluding entries lacking minimum taxonomic information. Creating and curating reference databases are labor-intensive tasks that require considerable time, proficiency with command-line tools, and the ability to optimize tool parameters. Several pipelines which are gene region agnostic, including OBITools3/ecoPCR (Boyer et al., 2016), RESCRIPt (Robeson et al., 2021), MetaCurator (Richardson et al., 2020), rCRUX and CRABS (Jeunen et al., 2023) have been developed to address these challenges and streamline the process.

The utility of plant DNA metabarcoding is dependent on accurately curated and regularly updated reference databases. Recognizing this need, in this study we generated and curated reference sequence databases for the most commonly sequenced *trnL* regions (CD, CH, and GH regions; **Figure 1**, adapted from Taberlet et al. (2007)) using tools with different methodological approaches: OBITools3/ecoPCR (*in silico* PCR), RESCRIPt (pairwise global alignment) and MetaCurator (hidden Markov model), and evaluated their classification performance.

**Figure 1.**
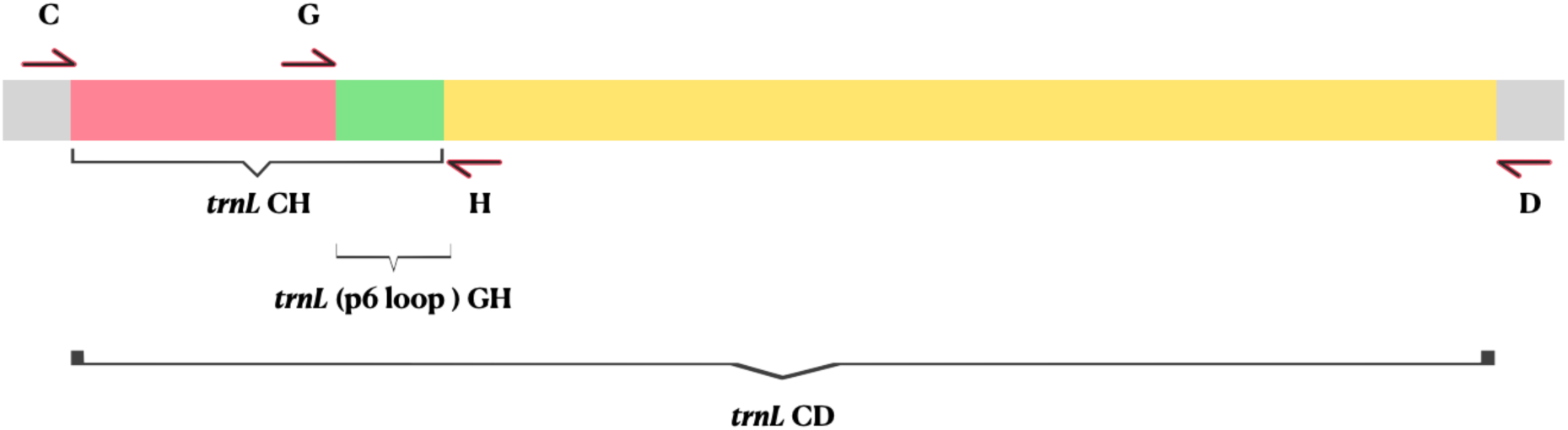
Primer positions and amplified regions of the chloroplast *trnL (UAA)* gene, adapted from (Taberlet et al., 2007). The figure illustrates the chloroplast *trnL* (UAA) intron, showing the positions of forward primers (**C and G**) and reverse primers (**H and D**), along with their corresponding amplified regions. All regions are proportionally scaled according to the *trnL* intron length in *Nicotiana tabacum*.

## 2. Materials and Methods

We evaluated three commonly used reference sequence database curation tools—OBITools3/ecoPCR, RESCRIPt, and MetaCurator—to construct *trnL* reference sequence databases **(Figure 2.A-C, detailed versions available in Supplementary Figure 1.A-B)**. We focused on primer pairs CD, CH, and GH (Taberlet et al., 2007): C 5′-CGAAATCGGTAGACGCTACG-3′, D 5′-GGGGATAGAGGGACTTGAAC-3′, G 5′-GGGCAATCCTGAGCCAA-3′, H 5′-CCATTGAGTCTCTGCACCTATC-3′; empirically determined sizes were ∼784–946 bp (CD), ∼100–500 bp (CH), and ∼10–143 bp (GH; P6 loop). A Google Scholar keyword search of the sequences of primers C and G (forward) and H and D (reverse) combinations revealed that a) 45% of studies used the GH primer pair targeting the highly conserved catalytic portion of the short P6 loop, b) 23% used CH, and c) another 23% used CD. Given only 9% used GD, this region was excluded from sequence curation in this study.

**Figure 2.**
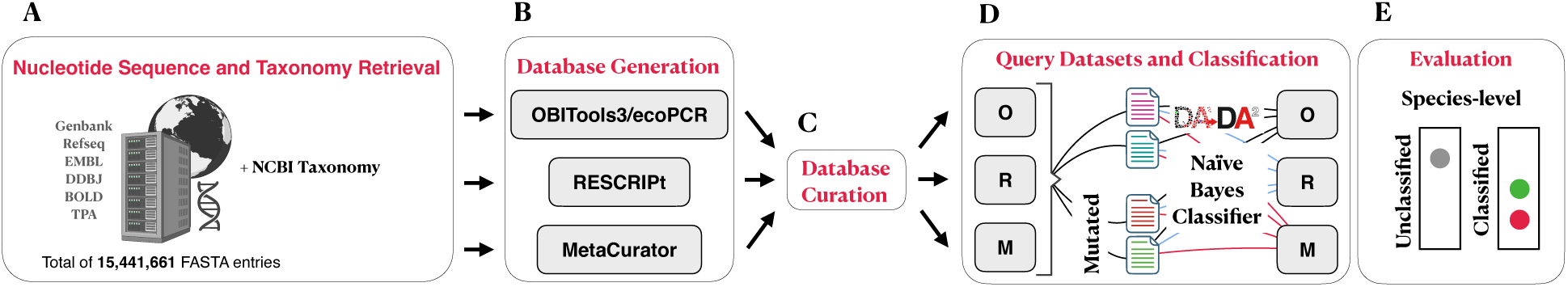
Workflow for *trnL* database generation, curation, and evaluation. Reference sequence databases for each trnL region were generated using OBITools3/ecoPCR (**O**), RESCRIPt (**R**), and MetaCurator (**M**). (**A, B**) Nucleotide sequences were retrieved from public repositories, and OBITools3/ecoPCR was used to generate reference sequences. These sequences were then used as seed sequences, restricted to plant taxa present in the US, for MetaCurator and RESCRIPt. (**C**) Raw sequences from the database generation step were curated using a standardized pipeline applied consistently across all tools. (**D**) Four query sets restricted to vascular plants were constructed to capture likely wind-pollinated and palynologically identifiable taxa in the US Taxonomic classification was performed using the Naïve Bayesian Classifier implemented in DADA2. (**E**) Classification performance was evaluated at the species levels using fraction classified, precision, and recall, based on correctly classified, misclassified, and unclassified results.

Software dependencies were managed and installed using Miniconda (Anaconda Inc., 2025). Unless otherwise specified, all processes, including nucleotide sequence acquisition and storage, script execution, data refinement, curation, and database assessment, were performed on the North Carolina State University Bioinformatics Research Center (BRC) Compute Cluster. Plant taxonomy information for species found in the United States (US) was sourced from the Natural Resources Conservation Service, U.S. Department of Agriculture (U.S. Department of Agriculture, 2016).

RESCRIPt was used as a QIIME2 plugin (qiime2-amplicon-2024.5.1). MetaCurator (v1.0.1) and its dependencies were installed on the BRC cluster following instructions from their respective GitHub pages (Richardson, 2019). OBITools3 (v3.0.1b26) was installed using the command `pip install OBITools3` (Python v3.7.12). CRABS (v0.1.8) used solely for sequence retrieval, was installed according to instructions on its GitHub page (Jeunen, 2022).

### 2.1 Nucleotide Sequence Retrieval

CRABS facilitates downloading sequencing data from online repositories, such as the NCBI via the `db_download` function. A total of 15,441,661 open-access, annotated Viridiplantae nucleotide sequences were retrieved from National Institutes of Health (NIH) GenBank, NCBI Reference Sequence Database (RefSeq), DDBJ (DNA Data Bank of Japan), European Molecular Biology Laboratory (EMBL) Nucleotide Sequence Database (EMBL-Bank), Third Party Annotation (TPA), all of which fall under the International Nucleotide Sequence Database Collaboration (INSDC: (Cochrane et al., 2011)). In addition, sequences were also obtained from the Barcode of Life Data System (BOLD: (Ratnasingham et al., 2024). Nucleotide sequences, excluding those from BOLD, were obtained from NCBI using specific keywords **(Supplementary Table 1).** A small number of FASTA entries, owing to NCBI server errors, were manually re-downloaded using accession numbers missing from the initial downloads.

### 2.2 Taxonomic Information Retrieval and Formatting

The FASTA headers of the downloaded files include accession numbers with versions, followed by sequence descriptions. Taxonomic information at the ranks of kingdom, phylum, class, order, family, genus, and species was extracted using these accession numbers via a Python script executed in Anaconda/Spyder (Conda v24.11.2, Spyder v5.5.1, Python v3.11.7). The script integrates the ETE Toolkit’s ncbi_taxonomy module (v3.1.3) and BioPython’s Entrez module (v1.83). The NCBI taxonomy database, downloaded on July 27, 2024, via `ncbi.update_taxonomy_database()`, was applied uniformly across all accession numbers. A subset of accession numbers lacked complete taxonomic information due to server timeouts and errors, requiring manual retrieval using additional scripts in Python and RStudio (v2023.6.1.524; (Posit Team, 2025)). In the final taxonomy file, all symbols were removed, and species names were formatted binomially, except for plant hybrids, which followed a format such as “*Citrus* × *latifolia*”.

The USDA PLANTS database, which contains identifiers, scientific names, and common names for each species (e.g., ABPR3, *Abrus precatorius* L., rosarypea respectively), was downloaded for all US states in August 2024. Binomial scientific names were then used with the August, 2, 2024 taxdump file to retrieve taxIDs and full taxonomic rank information via the taxizedb (v0.3.1; (Chamberlain et al., 2023)) library in RStudio. Approximately 18% of scientific names did not yield a taxID and therefore these entries were removed.

### 2.3 Seed Sequence Generation for RESCRIPt and MetaCurator

OBITools3/ecoPCR was employed for seed sequence generation. All downloaded sequences and associated NCBI taxonomy files were imported as obidms (data management system) objects in this format: >Accession number.{version number} TAXID={taxID}; species_name={binary name} (e.g., >NC_001879.2 TAXID=4097; species_name=Nicotiana tabacum).

After importing the files, ecoPCR was executed to generate *in silico* fragments for the *trnL* CD, CH, and GH regions. The corresponding primer pairs were used with the following parameters: `-e 3` (allowing up to three mismatches per primer), `-l 1` (setting a minimum amplified fragment length, excluding primers), and `-L 5000` (setting a maximum amplified fragment length, excluding primers). After completing the ecoPCR step, the resulting obidms objects were exported as FASTA files for each *trnL* region. FASTA entries were split into species-specific files, and CD-HIT (v4.8.1; (Li & Godzik, 2006)) was run with exact match parameters `-c 1.0 -n 10 -T 0 -d 0` to generate .clstr files, which were parsed with a Python script to retain shorter fragments instead of CD-HIT’s default longer sequences.

Seed sequences were filtered to include only species present in the USDA plant taxonomy dataset using taxID information, yielding 15,334, 16,423, and 43,458 entries for the *trnL* CD, CH, and GH regions, respectively. All FASTA files were then split by genus, and the number of seed sequences was reduced to prevent slowdowns in downstream processes; a single representative sequence was randomly selected for each unique genus, followed by an additional CD-HIT run to remove exact duplicates. This process yielded 1,380, 1,174, and 1,720 entries for the *trnL* CD, CH, and GH regions, respectively, which were then used as seed sequences for RESCRIPt and MetaCurator.

### 2.4 Database Generation

**2.4.1 OBITools3/ecoPCR**

We followed the process described in Section 2.3*: Seed Sequence Generation for RESCRIPt and MetaCurator.* Briefly, all formatted input FASTA files and the NCBI taxonomy file were imported into obidms, and the ecoPCR module was executed to generate *in silico* PCR fragments for each *trnL* region. The raw FASTA outputs were exported from obidms yielding three raw FASTA files—one corresponding to each *trnL* region.

#### 2.4.2 RESCRIPt

Once the input sequences, seed sequences and taxonomy file were imported into QIIME2, the qiime rescript extract-seq-segments command was executed with --p-perc-identity 0.8 and --p-min-seq-len ${min_seq_len}, set ${min_seq_len} to the corresponding minimum sequence length for each *trnL* region, parameters.

#### 2.4.3 MetaCurator

All FASTA files were split into ∼1-2 gigabyte subsets using the seqkit command-line tool (Shen et al., 2024) with parameters `split2 --by-part {size of the file}`. This step was implemented to expedite processing due to the large size of the combined input sequences. Corresponding taxonomy files were then created for each subset using a Python script. The `MetaCurator.py` script was subsequently executed for each subset with parameters as suggested in the github workflow `-is 6,6,3,3,3`; `-cs 1.0,0.98,0.95,0.9,0.85`; `-e 0.005`; `-ct True`; `-tf True`. Here, `-is` specifies the number of searches to perform during each round of extraction, `-cs` defines the minimum HMM coverage per round for an extracted amplicon reference sequence, `- e` sets the E-value threshold for HMM searches, `-ct` enables cleaning of taxonomic lineages to remove common NCBI artifacts, and `-tf` indicates that the taxonomy file is in Taxonomizr format.

### 2.5 Database Curation

Raw database files for each of the three methods (OBITools3/ecoPCR, RESCRIPt and MetaCurator) underwent the same curation steps **(Figure 2.C, detailed versions available in Supplementary Figure 1.B)** to ensure consistency. Briefly, four steps were completed to curate the sequences:

**1.** Filtering ambiguous sequences: All sequences containing any ambiguous characters (anything other than A, T, C, or G) were removed.
**2.** Filtering by Taxonomy: Sequences missing both genus and species taxonomic information were excluded, along with entries containing specific keywords such as “nan prasinophyte” or “nan Streptophyta”. In addition, given the focus of this study, green algae sequences were also excluded from the final databases by removing entries where the phylum-level taxonomy indicated Prasinodermophyta or Chlorophyta.
**3.** Filtering by sequence length: Based on examination of unfiltered land plant sequences generated by RESCRIPt and OBITools3/ecoPCR, acceptable size ranges were determined as follows: *trnL*-CD: 234-1566 nt, *trnL*-CH: 81-484 nt, and *trnL*-GH: 8-220 nt.
**4.** Dereplicate sequences: Multiple entries for the same species introduce redundancy in any database. To address this, FASTA entries were first split by species, and CD-HIT was applied `cd-hit-est -i “$input_file” -o “$output_file” -c 1.0 -n 10 -T 0 -d 0` to remove exact duplicate sequences within each species-specific FASTA file. By default, CD-HIT clusters identical sequences and removed the shortest ones.

### 2.6 Classification

We used the Naïve Bayesian Classifier (Wang et al., 2007) implemented in the DADA2 package (Callahan et al., 2016) for taxonomic classification, applying default parameters using the command: `taxa <- assignTaxonomy(${query fasta file}, ${database fasta file}, multithread = TRUE)`.

### 2.7 Evaluation

#### 2.7.1 Simulated query datasets

Simulated query datasets were composed of vascular plants, including Angiosperms (flowering plants) and Gymnosperms (seed-producing plants) found in the US. The selection was intended to capture possible wind-pollinated (anemophilous) taxa and those that can be identified palynologically from surface soil samples collected in the US. Four simulated test sets were created (**Figure 2.D**; detailed version available in **Supplementary Figure 1.C-D**): **(1)** Random Combination (3,000 entries), consisting of an equal number of randomly selected entries from OBITools3/ecoPCR, RESCRIPt and MetaCurator databases; **(2)** Common Species (3,000 entries), composed of randomly selected entries from species shared across all three databases; **(3)** Mutated Random Combination, and **(4)** Mutated Common Species, where random mutations were introduced while ensuring that mutations did not exceed the species-wise average alignment percentage (96.52%, based on sequences generated by OBITools3/ecoPCR). The Random Combination query set was designed to evaluate how well each database classifies sequences curated both by itself and by other tools, highlighting the impact of curation differences of the tools.

#### 2.7.2 Evaluation of Classification

Classification was evaluated (**Figure 2.E**; detailed version available in **Supplementary Figure 1.E**) at family, genus and species levels. Species-level evaluations are presented in the main manuscript, and comparative results for the family and genus ranks are detailed in the supplementary materials. If a sequence was correctly assigned at a given taxonomic level, it was counted as **“correctly classified”.** Incorrect assignments were categorized as **“misclassified”**, while cases where no classification was made at that level were recorded as **“unclassified”**. Additionally, we reported commonly used performance metrics, including Fraction Classified, Precision and Recall (Performance metrics definitions and calculations are in **Table 1**).

**Table 1.** Calculation of Performance Metrics for Database Taxonomic Classification. The Fraction classified (**FC**) represents the proportion of classified entries among all entries (**N**). Precision (**P**) measures the proportion of correctly classified entries among those classified. Recall (**R**), defined as the product of FC and P, quantifies the proportion of correctly classified entries among all entries (**N**).

| Metrics | Calculation |
| --- | --- |
| **Fraction classified (FC)** | (Correctly Classified + Misclassified) / N |
| **Precision (P)** | Correctly Classified / (Correctly Classified + Misclassified) |
| **Recall (R)** | Correctly Classified / N |

#### 2.7.3 Assessment of Execution Time and Peak Memory Usage

The GenBank-1 file, consisting of 1,058,168 FASTA entries, was used to evaluate total execution time (minutes) and peak memory usage (gigabytes) across all tools and *trnL* regions. Each job was submitted using the `sbatch` SLURM command with `-c 12` (number of CPU cores, enabling multi-threading and parallel processing) and `--mem=100G` (total memory (RAM) allocated per node).

Execution time and memory usage for OBITools3/ecoPCR were assessed during the import of sequence and taxonomy files into the DMS, execution of ecoPCR, and export of the DMS file to FASTA format. In the case of RESCRIPt, input sequence and taxonomy files were pre-converted into QIIME 2 artifacts, and performance evaluation focused solely on the `extract-seq-segments` command, which represented the primary computational bottleneck. MetaCurator’s execution time and memory usage were measured during the run of the `MetaCurator.py` script, which automated the entire workflow. SLURM logs were generated separately for each job, capturing total execution time (default: seconds) and peak memory usage (default: kilobytes), which were then converted to minutes (min) and gigabytes (GB) for reporting.

## 3. Results

Three methodologically distinct database curation tools—MetaCurator, RESCRIPt, and OBITools3/ecoPCR—were used to generate reference sequence databases for *trnL* regions (CD, CH, and GH). A common initial curation step included ambiguous (degenerate) base removal, taxonomy cleanup, length-based filtering, and sequence dereplication. The number of raw FASTA entries and the remaining entries after each curation step for all *trnL* regions across tools are provided in **Supplementary Table 2**. Ambiguous sequences accounted for no more than 2% of sequences in OBITools3/ecoPCR and 5% in MetaCurator but RESCRIPt initially generated millions of sequences containing ambiguous sequences in the raw file. **Figure 3** shows the sequence length distribution for both raw and curated databases across tools for all three *trnL* regions.

**Figure 3.**
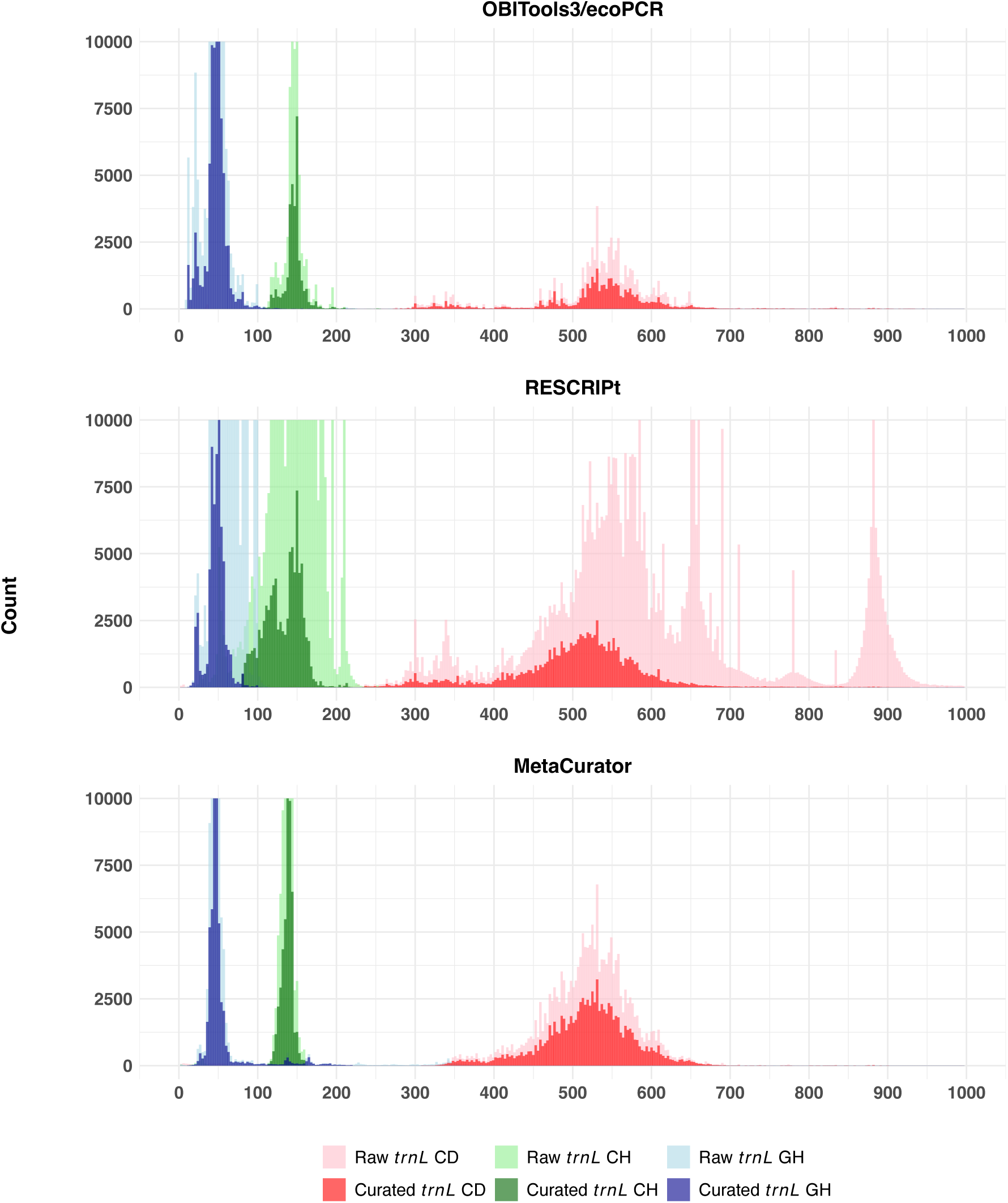
Raw and curated *trnL* sequence length distributions across curation tools. Histogram plots showing the distribution of raw and curated *trnL* sequence lengths for OBITools3/ecoPCR, RESCRIPt, and MetaCurator. Each color represents a *trnL* region (CD, red; CH, green; GH, blue), with light shades indicating raw sequences and dark shades indicating curated sequences. The y-axis limit was set to 10,000 counts to visualize lower-count bins, while the x-axis limit was set to 1,000 nucleotides, as the number of entries beyond that point was minimal.

### 3.1 Breadth of Taxonomy Recovery

Each tool showed variation in the breadth of taxonomy recovery, as measured by the number of unique taxonomic entries retained across the *trnL* regions (**Table 2**, species-level Venn diagram is in **Figure 4**; family- and genus-level visualizations are available in **Supplementary Figure 2-3**, respectively, and the comprehensive taxonomic comparison table is provided in **Supplementary Table 3**). In terms of species retrieval, MetaCurator recovered the highest number of unique species in the *trnL* CD region, RESCRIPt in the *trnL* CH region, and OBITools3/ecoPCR in the *trnL* GH region.

**Figure 4.**
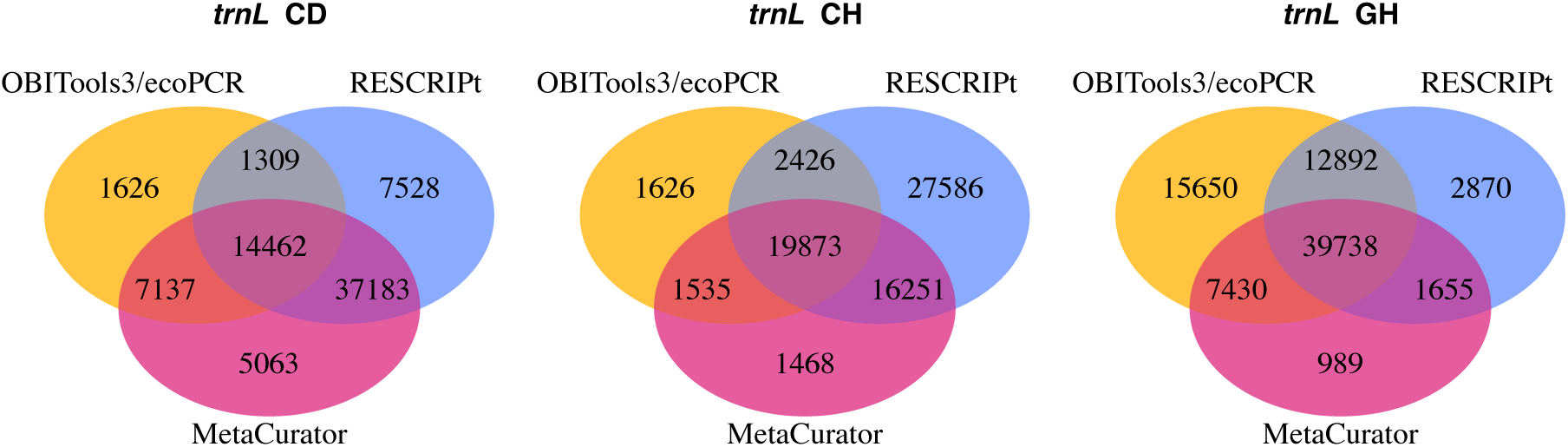
Number of unique species retained by each curation tool for each *trnL* region. Venn diagrams illustrating the number of unique and shared species generated by OBITools3/ecoPCR (yellow), RESCRIPt (blue), and MetaCurator (pink) for *trnL* CD, CH, and GH regions.

**Table 2.**
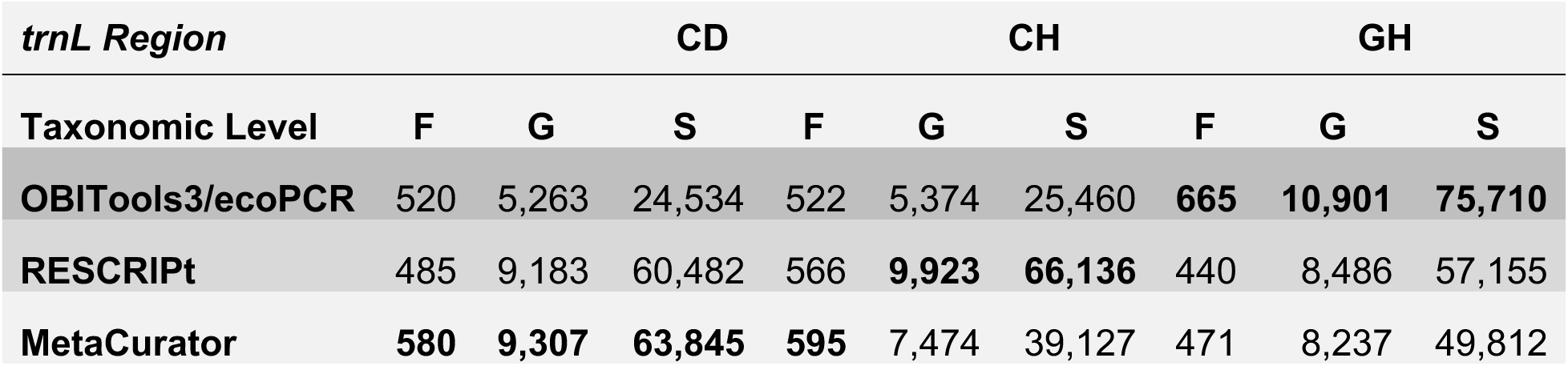
Number of unique families (F), genera (G), and species (S) retained by each curation tool across all *trnL* regions. Bold values indicate the highest count among curation tools.

| *trnL* Region | CD |  |  | CH |  |  | GH |  |  |
| --- | --- | --- | --- | --- | --- | --- | --- | --- | --- |
| Taxonomic Level | F | G | S | F | G | S | F | G | S |
| OBITools3/ecoPCR | 520 | 5,263 | 24,534 | 522 | 5,374 | 25,460 | **665** | **10,901** | **75,710** |
| RESCRIPT | 485 | 9,183 | 60,482 | 566 | **9,923** | **66,136** | 440 | 8,486 | 57,155 |
| MetaCurator | **580** | **9,307** | **63,845** | **595** | 7,474 | 39,127 | 471 | 8,237 | 49,812 |

### 3.2. Classification Evaluation

We evaluated the reference databases using classification-outcome metrics (Correctly Classified (C), Misclassified (M), Unclassified(U)) and performance metrics—Fraction Classified (FC), Precision (P), and Recall (R). Analyses were performed at the family, genus and species levels across four simulated query datasets: Random Combination, Common Species, Mutated Random Combination and Mutated Common Species. Here we concentrate on species-level results for the mutated query datasets (Mutated Random Combination and Mutated Common Species), which most closely reflects real-world scenario. Comparative performance metrics across mutated query datasets and *trnL* regions is presented in **Table 3** and **Figure 5**.

**Figure 5.**
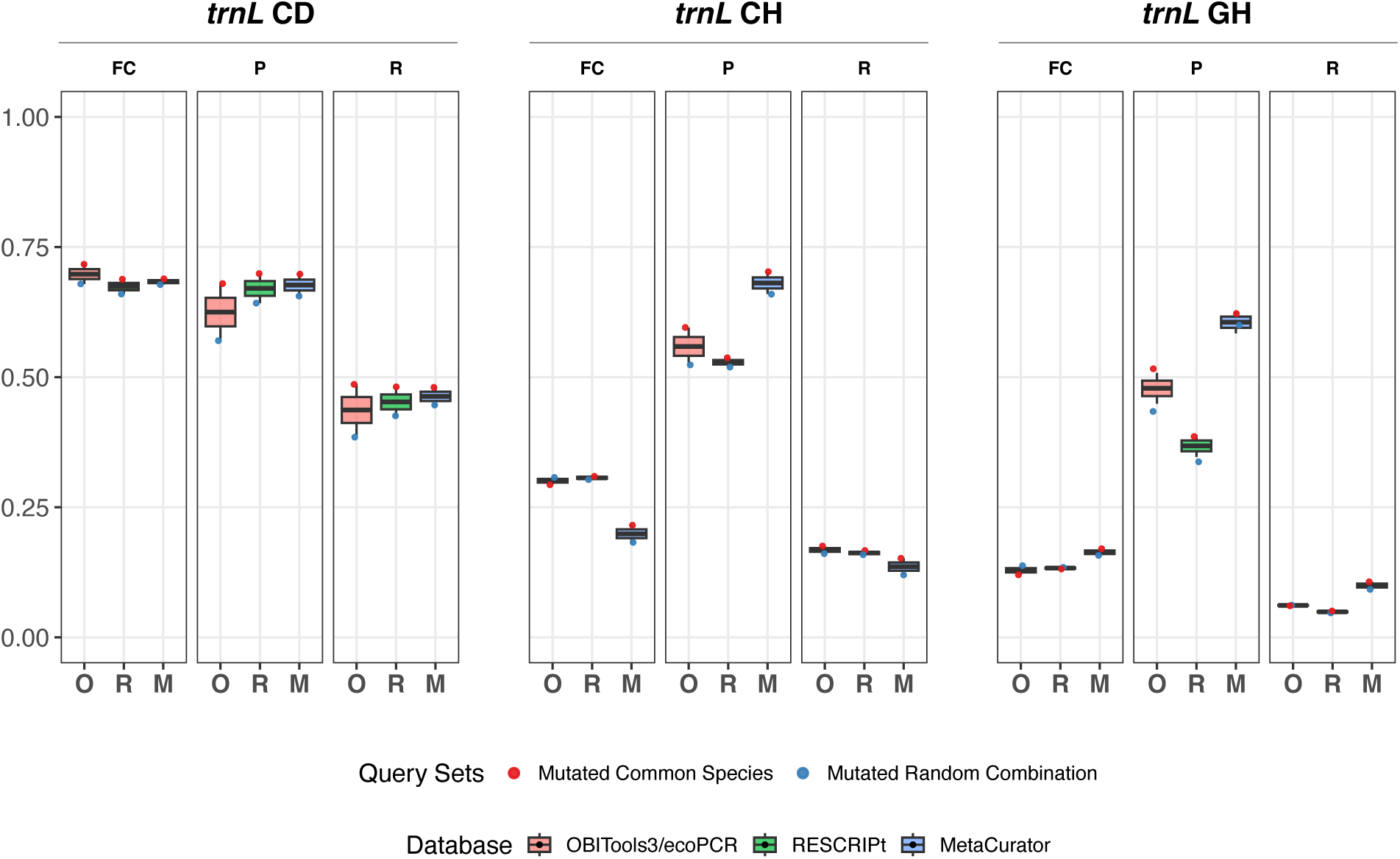
Performance metrics across *trnL* CD, CH, and GH regions for each tool at the species level. Boxplots show the fraction classified (**FC**), precision (**CP**), and recall (**R**) for OBITools3/ecoPCR (**O**), RESCRIPt (**R**), and MetaCurator (**M**) reference databases across the *trnL* CD, CH, and GH regions. Datapoints are shown for both Mutated Common Species and Mutated Random Combination query sets.

**Table 3.** Taxonomic classification metrics across mutated query sets and all databases. Comparison of taxonomic classification performance at the species levels across mutated random combination (**MRC**), and mutated common species (**MCS**) query sets and all databases. Metrics include the number of correctly classified (**C**), unclassified (**U**), and misclassified (**M**) entries, along with fraction classified (**FC**) and precision (**CP**) and recall (**R**). Bold values indicate the highest performance for each metric.

| Query | Databases | *trnL* CD |  |  |  |  |  | *trnL* CH |  |  |  |  |  | *trnL* GH |  |  |  |  |  |
| --- | --- | --- | --- | --- | --- | --- | --- | --- | --- | --- | --- | --- | --- | --- | --- | --- | --- | --- | --- |
|  |  | C | U | M | FC | P | R | C | U | M | FC | P | R | C | U | M | FC | P | R |
| **MCS** | **OBITools3/ecoPCR** | **1,461** | **850** | 689 | **0.717** | 0.680 | **0.487** | **524** | 2,119 | 357 | 0.294 | 0.595 | **0.175** | 184 | 2,638 | **178** | 0.121 | 0.508 | 0.061 |
| **MCS** | **RESCRIPT** | 1,443 | 937 | **620** | 0.688 | **0.699** | 0.481 | 499 | **2,071** | 430 | **0.310** | 0.537 | 0.166 | 153 | 2,607 | 240 | 0.131 | 0.389 | 0.051 |
| **MCS** | **MetaCurator** | 1,442 | 933 | 625 | 0.689 | 0.698 | 0.481 | 455 | 2,352 | **193** | 0.216 | **0.702** | 0.152 | **320** | **2,490** | 190 | **0.170** | **0.627** | **0.107** |
| **MRC** | **OBITools3/ecoPCR** | 1,161 | **963** | 876 | **0.679** | 0.570 | 0.387 | **484** | **2,075** | 441 | **0.308** | 0.523 | **0.161** | 185 | 2,588 | 227 | 0.137 | 0.449 | 0.062 |
| **MRC** | **RESCRIPT** | 1,271 | 1,020 | 709 | 0.660 | 0.642 | 0.424 | 473 | 2,090 | 437 | 0.303 | 0.520 | 0.158 | 140 | 2,596 | 264 | 0.135 | 0.347 | 0.047 |
| **MRC** | **MetaCurator** | **1,334** | 967 | **699** | 0.678 | **0.656** | **0.445** | 361 | 2,453 | **186** | 0.182 | **0.660** | 0.120 | **275** | **2,529** | **196** | **0.157** | **0.584** | **0.092** |

Although species-level results are emphasized, analogous trends were observed for the mutated datasets at the genus and family levels, with inter-database contrasts becoming less pronounced at higher taxonomic ranks (**Supplementary Figure 4**). This pattern generally held when extended to all query datasets, although a few minor deviations were present (**Supplementary Figure 5**).

In the *trnL* CD region, classifications obtained using the OBITools3/ecoPCR reference database produced a slightly greater number of classified entries, but this increase was accompanied by a higher rate of misclassification. Consequently, classifications generated using MetaCurator and RESCRIPt yielded higher precision and recall.

In the *trnL* CH region, the use of the OBITools3/ecoPCR and RESCRIPt reference databases resulted in markedly more classified entries than MetaCurator. Yet although overall recall was marginally higher for those two databases, their elevated misclassification rates resulted in MetaCurator attaining the greatest precision.

In the *trnL* GH region, classifications backed by MetaCurator consistently outperformed those using OBITools3/ecoPCR or RESCRIPt across all performance metrics.

### 3.3 Evaluation of Execution Time and Peak Memory Usage

Execution time and peak memory usage are detailed in **Table 4**. OBITools3/ecoPCR was the fastest, with execution time not exceeding 14 minutes and peak memory usage remaining below 16 GB when processing the GenBank-1 file, which contains 1,058,168 FASTA entries. RESCRIPt ran slower than OBITools3/ecoPCR but outperformed MetaCurator in execution time, with the fastest processing observed for the *trnL* GH region, followed by *trnL* CH and *trnL* CD. Notably, the peak memory usage of RESCRIPt followed an inverse trend to execution time, being lowest for *trnL* CD, followed by *trnL* CH and *trnL* GH. MetaCurator is optimized for memory usage (not exceeding 17 GB) but execution time varied by region, being shortest for *trnL* CH, followed by *trnL* GH, and longest for *trnL* CD.

**Table 4.** Execution time and peak memory usage across tools for each *trnL* region. Total execution time and peak memory usage for processing *trnL* CD, CH, and GH regions using OBITools3/ecoPCR (**O**), RESCRIPt (**R**), and MetaCurator (**M**) for GenBank-1 file (1,058,168 FASTA entries). Bold values indicate the highest performance for each metric.

|  | Time and Space | O | R | M |
| --- | --- | --- | --- | --- |
| ***trnL* CD** | Total execution time (minutes) | **13.95** | 1,685.58 | 9,109.15 |
|  | Peak memory usage (Gigabyte) | **15.92** | 60.83 | 16.51 |
| ***trnL* CH** | Total execution time (minutes) | **12.40** | 238.50 | 486.67 |
|  | Peak memory usage (Gigabyte) | **15.92** | 77.41 | 16.60 |
| ***trnL* GH** | Total execution time (minutes) | **11.78** | 62.35 | 1,361.63 |
|  | Peak memory usage (Gigabyte) | **15.92** | 83.11 | 16.60 |

## 4. Discussion

This study evaluated reference databases generated for the *trnL* CD, CH, and GH regions using three curation tools—OBITools3/ecoPCR, RESCRIPt, and MetaCurator—each implementing a distinct methodological framework. Computational demands are often a limiting factor in database generation from large sequence inputs, and execution metrics from **Table 4** show that OBITools3/ecoPCR requires the least computational resources and provides uniform performance across *trnL* regions. This low computational demand arises from its *in-silico* PCR approach, which relies on the Agrep pattern-matching algorithm (Ficetola et al., 2010). MetaCurator exhibits similarly consistent memory usage but requires the longest runtime due to its iterative search algorithm, whereas RESCRIPt consumes more memory because of its pairwise global alignment algorithm and yields intermediate runtimes.

Although OBITools3/ecoPCR is computationally efficient, it requires both primer sites to be present in input sequences. Many INSDC submissions may lack one or both primer-binding regions, which reduces the number of sequences retained. We observe the effect of this pattern in our results; OBITools3/ecoPCR retains the smallest number of genera and species for the CD and CH regions (**Table 2**). The GH region, however, represents an exception: OBITools3/ecoPCR retrieves more entries and unique taxa, likely due to the higher frequency of intact GH primer regions in the input sequences. Another explanation could be that, as previously reported, GH primers are highly conserved (Taberlet et al., 2007), which increases the likelihood that sequences pass the mismatch-threshold filter `-e 3` used by OBITools3/ecoPCR.

Despite the larger *trnL* GH database retrieved by OBITools3/ecoPCR, MetaCurator outperforms all tools in every comparative metric across taxonomic levels (**Supplementary Table 6**). This discrepancy may reflect the limited resolving power of the GH region, the shortest among the *trnL* markers. In addition, taxonomic classification was performed using the Naïve Bayesian Classifier, one of the most widely used tools for taxonomically classifying marker-gene sequences. Several implementations are available: in this study the implementation in the DADA2 R package with default parameters was used. However, the *trnL* GH region includes particularly short sequences (typical range of 8-220 bp), and the DADA2 classifier imposes minimum length requirements—20 nucleotides for reference sequences and 50 nucleotides for input sequences—for reliable taxonomic assignment. As a result, many sequences in the *trnL* GH dataset remained unclassified, even at the family level, along with the low resolution of the region. Unclassification rates were 50% and 40% for OBITools3/ecoPCR and MetaCurator, respectively. The low resolution and high number of highly similar sequences increase the frequency of classification ties, producing elevated unclassification rates. These findings suggest that users may require additional refinement of databases by restricting taxonomic scope to lineages relevant to their study rather than relying on an unfiltered global database. RESCRIPt, on the other hand, shows the lowest precision, likely due to false-positive recruitment driven by its pairwise global alignment strategy, which can admit spurious matches and allow boundary drift for short *trnL* GH sequences, especially when combined with short seed sequences and the similarity parameter we applied. We further examined RESCRIPt’s behavior for the *trnL* GH region by comparing the lengths of identical input sequences (matched by accession number) returned by RESCRIPt and MetaCurator against those produced by OBITools3/ecoPCR (**Supplementary Figure 6**). We found that RESCRIPt exhibited larger sequence-length differences for the *trnL* GH region than for the other regions. This additional pattern lends modest support to our interpretation of RESCRIPt’s performance in this region.

Even though RESCRIPt and MetaCurator are more computationally intensive, they do not depend on primer presence and therefore retain more sequences and a broader set of unique taxa for the CD and CH regions. Both methods, however, rely on seed sequences to recruit homologous records, and poorly chosen seeds with limited diversity may constrain taxonomic breadth and introduce representation bias. Based on the collective results, in *trnL* CD—the longest region—RESCRIPt and MetaCurator show comparable performance, with both achieving higher precision and recall than OBITools3/ecoPCR, likely owing to the absence of primer-binding sites in a substantial subset of sequences. Given these observations, either RESCRIPt or MetaCurator is suitable for constructing *trnL* CD databases (Classification metrics are detailed in **Supplementary Table 4**).

For *trnL* CH, RESCRIPt maintains strong performance in terms of the total number of retrieved entries; however, the number of unique species retrieved exclusively by RESCRIPt is markedly higher than that obtained by the other tools (**Figure 4**). Given the CH region also has limited taxonomic resolution compared to *trnL* CD region, we observed a corresponding decline in precision for RESCRIPt, likely reflecting low intraspecific variability in this region. Taken together, there is no single tool performs better across all evaluation metrics for the *trnL* CH region: MetaCurator yields the fewest misclassified entries, yet its fraction of classified sequences is low, which constrains its practical utility. Users should therefore consult **Supplementary Table 5** and **Supplementary Figures 4-5** to determine which combination of tool and taxonomic level aligns best with the needs of their specific application.

Beyond algorithmic and performance differences, database construction also presents practical challenges. Preparing taxonomy and sequence files to match tool-specific input formats often requires extensive preprocessing. All tools required some degree of input modification; however, none was more demanding than the others. This challenge persists because INSDC data structures continue to evolve, and we are hoping that emerging generative AI utilities will help users adapt input files to the changing requirements of these tools.

This study has limitations that should be considered when evaluating the findings, as detailed in the following points:

**a)** Uniform parameter configuration across *trnL* regions: Database curation parameters were applied uniformly across all *trnL* regions rather than optimized individually. OBITools3/ecoPCR was executed with an error rate of three following the Wolf tutorial (Coissac, 2023), RESCRIPt parameters were adopted from the QIIME2 forum documentation (Robeson, 2022), and MetaCurator settings followed the GitHub workflow. Users may need to optimize these parameters using small test sets considering the *trnL* region of interest.
**b)** Seed sequence configuration for RESCRIPt and MetaCurator: Comparability across tools was maintained by initializing both RESCRIPt and MetaCurator with the same seed sequences generated by OBITools3/ecoPCR. While this uniform framework facilitated consistent benchmarking, it may have limited each tool’s performance, as optimal configurations likely differ depending on whether database completeness or computational efficiency is prioritized.
**c)** Splitting of input sequences: MetaCurator’s iterative HMM-based extraction updates the reference profile with each round, but partitioning the input data into smaller subsets may have disrupted this iterative refinement process and reduced database expansion efficiency.
**d)** Query datasets: Identifying an appropriate simulated dataset for performance evaluation remained challenging. The query sets generated from all three tools likely contained false positives and exhibited unbalanced taxonomic representation. This bias likely stems from uneven sequencing efforts across plant taxa, driven by differences in their economic and research significance. Furthermore, the emphasis on vascular plants in the US, particularly wind-pollinated taxa, amplified this imbalance and led to an overrepresentation of Poaceae family across all datasets. A suitable query set would ideally comprise synthetic samples with known taxonomic composition to provide a reliable benchmark, but such data were unavailable for this study.

In conclusion, each tool offers distinct strengths and limitations, and the choice depends on the *trnL* region and the user’s available resources. Considering species-level evaluations, RESCRIPt and MetaCurator performed strongly for the *trnL* CD region, OBITools3 and RESCRIPt showed comparable performance for the *trnL* CH region, and MetaCurator provided the best performance for the *trnL* GH region.

We also provide curated databases and taxonomy files for each *trnL* region and tool in plain and DADA2 compatible FASTA formats (https://doi.org/10.5281/zenodo.17969450). Although this study was designed around a specific taxonomic scope, the resulting *trnL* reference sequence databases are sufficiently comprehensive for broader applications. The methods and findings presented here can assist researchers in selecting appropriate tools based on the target marker region and study design. Users interested in utilizing these databases are advised to review the associated taxonomy files and assess the breadth of taxonomic coverage relevant to their plant taxa of interest. For those wishing to generate custom reference databases, the workflow outlined in this study provides a practical framework for building a single database or comparing multiple databases tailored to specific taxonomic groups.

## Acknowledgements

This research was sponsored by the Army Research Office and was accomplished under Cooperative Agreement Number W911NF-21-2-0129. This manuscript is a preprint and is currently under review. It has not been peer reviewed.

## Data Availability Statement

All reference databases, taxonomy, and evaluation files are available at Zenodo (https://doi.org/10.5281/zenodo.17969450). The complete computational workflow and scripts are available on GitHub (https://github.com/oskuddar/trnL_DB).

## Author Contributions

**O.S.K.** and **B.J.K.** conceptualized the study. Methodology, software, formal analysis, data curation, and writing of the original draft were performed by **O.S.K**. Validation was conducted by **O.S.K.** and **B.J.K**. Writing (review and editing) was a collaborative effort by **B.J.K.**, **K.A.M.**, and **O.S.K**. Supervision and funding acquisition were provided by **B.J.K.** and **K.A.M**. All authors have read and agreed to the published version of the manuscript.

## Disclosure

Code development for data processing and analysis was assisted by the generative AI tool ChatGPT (OpenAI). The conceptual framework, analytical design, and mathematical formulations were developed by the corresponding author, who provided explicit instructions for code generation. All AI-generated code was manually reviewed, modified where necessary, and validated by the corresponding author to ensure correctness and consistency with the intended methodology. The corresponding author takes full responsibility for the final code, analyses, and results.

The views and conclusions contained in this document are those of the authors and should not be interpreted as representing the official policies, either expressed or implied, of the Army Research Office or the US Government. The US Government is authorized to reproduce and distribute reprints for Government purposes, notwithstanding any copyright notation herein.

