## Supplementary Figures and Tables for "Generating, curating, and evaluating *trnL* reference sequence databases: Benchmarking OBITools3/ecoPCR, RESCRIPt, and MetaCurator"

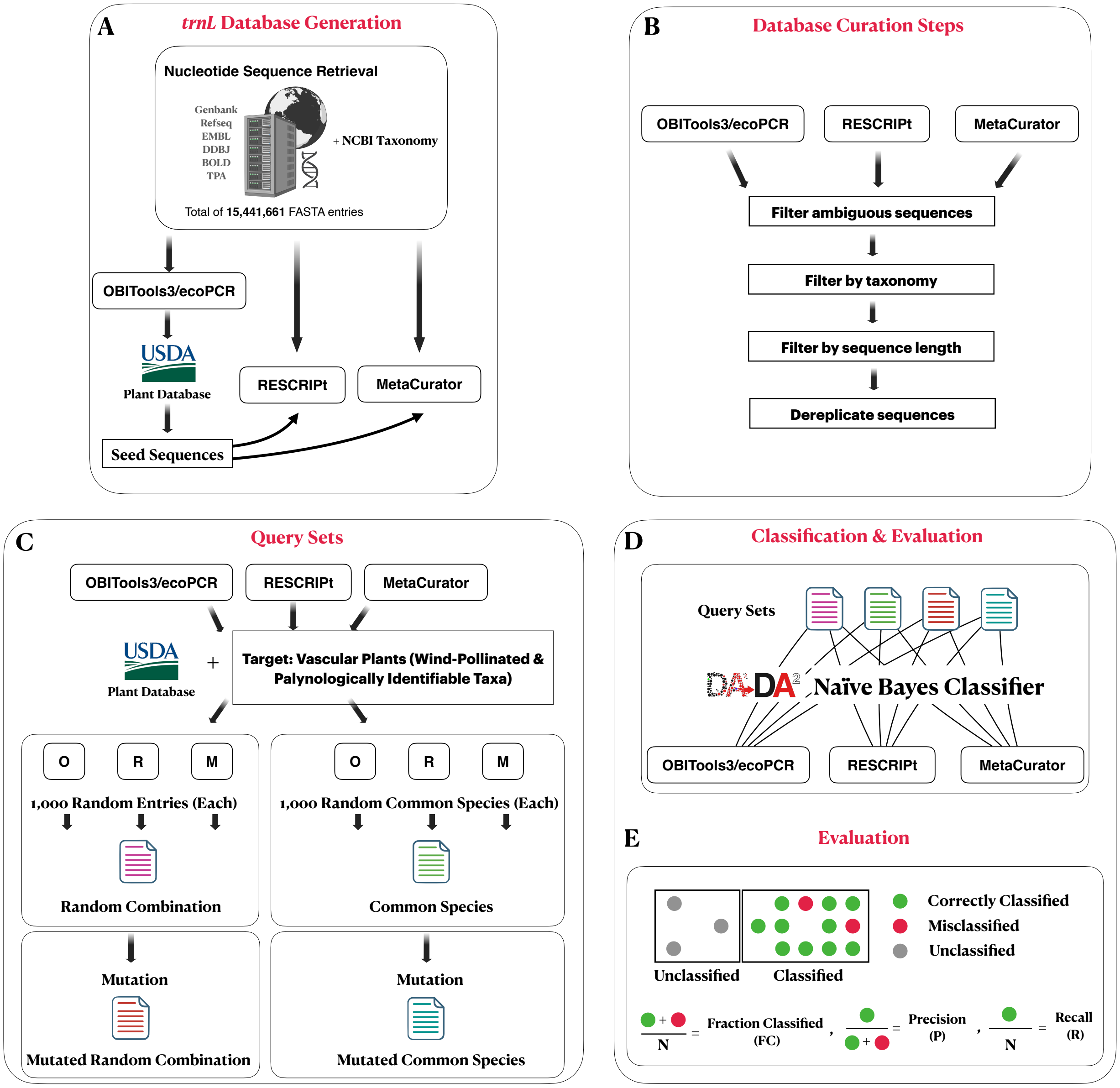


**Supplementary Figure 1. Detailed workflow for *trnL* database generation, curation, query construction, taxonomic classification and evaluation.**
Reference sequence databases were generated using OBITools3/ecoPCR (**O**), RESCRIPt (**R**), and MetaCurator (**M**), and downstream classification performance was evaluated.

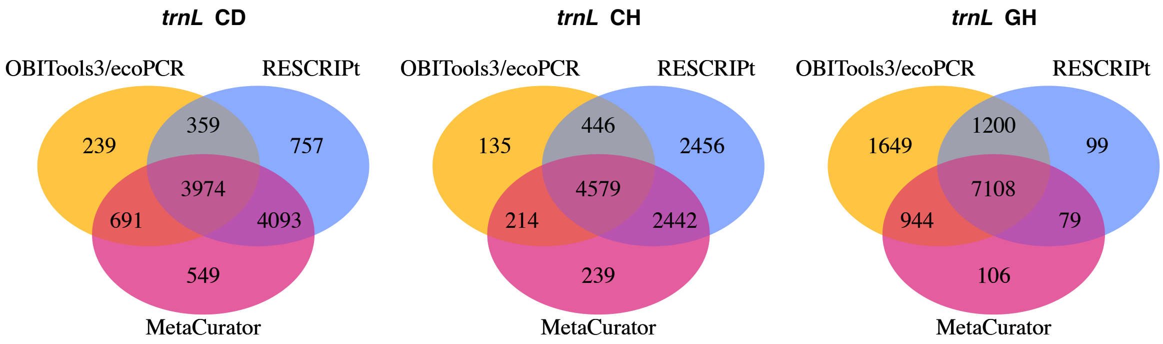


**Supplementary Figure 2. Number of unique genera retained by each curation tool for each *trnL* region.**

Venn diagrams illustrating the number of unique and shared genera generated by OBITools3/ecoPCR (yellow), RESCRIPt (blue), and MetaCurator (pink) for *trnL* CD, CH, and GH regions.

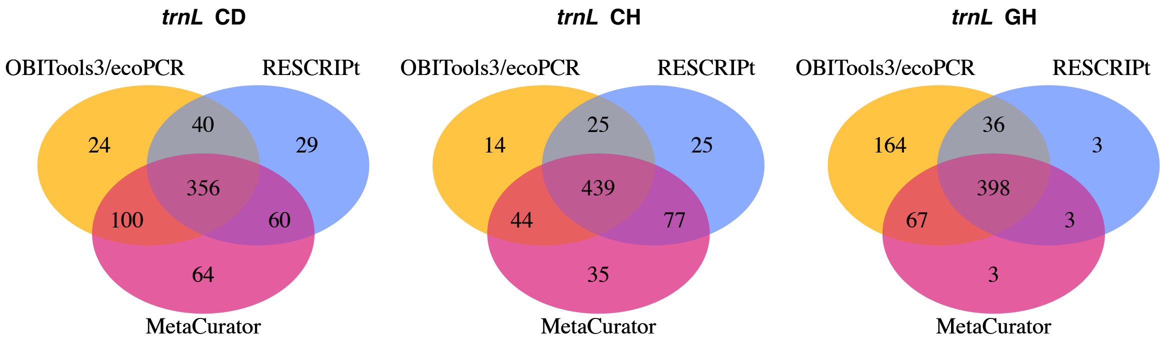


**Supplementary Figure 3. Number of unique families retained by each curation tool for each *trnL* region.**

Venn diagrams illustrating the number of unique and shared families generated by OBITools3/ecoPCR (yellow), RESCRIPt (blue), and MetaCurator (pink) for *trnL* CD, CH, and GH regions.

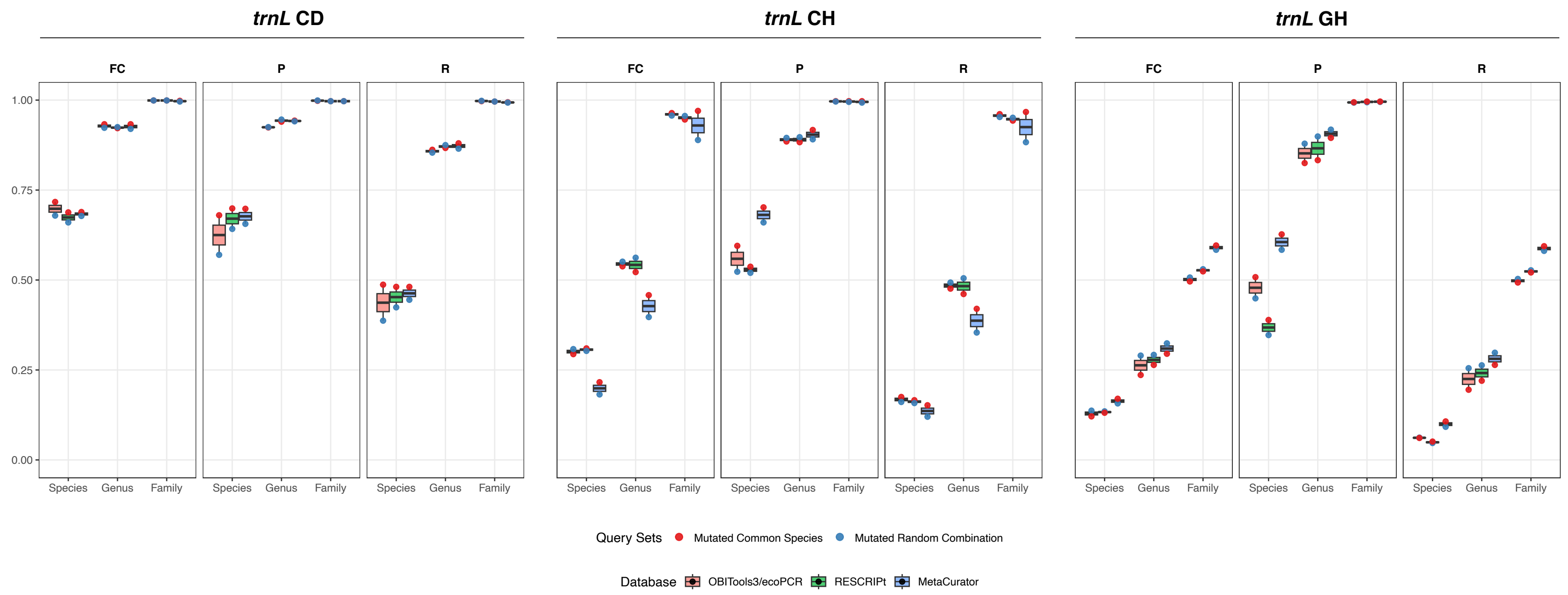


**Supplementary Figure 4. Performance metrics across *trnL* CD, CH, and GH regions for each tool at the family, genus, and species levels.**

Boxplots show the fraction classified (**FC**), precision (**CP**), and recall (**R**) for OBITools3/ecoPCR (**O**), RESCRIPt (**R**), and MetaCurator (**M**) reference databases across the *trnL* CD, CH, and GH regions for mutated query sets. Data points are shown for both Mutated Common Species and Mutated Random Combination query sets.

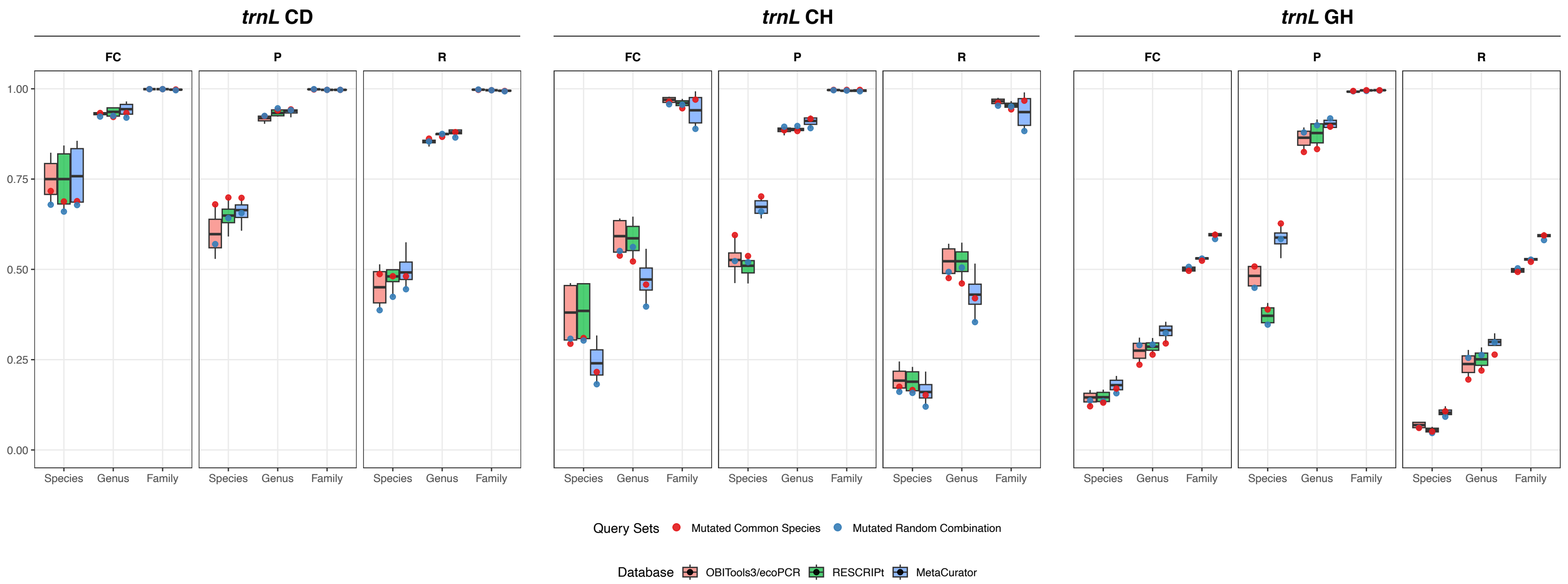


**Supplementary Figure 5. Performance metrics across *trnL* CD, CH, and GH regions for each tool at the family, genus, and species levels.**

Boxplots show the fraction classified (**FC**), precision (**CP**), and recall (**R**) for OBITools3/ecoPCR (**O**), RESCRIPt (**R**), and MetaCurator (**M**) reference databases across the *trnL* CD, CH, and GH regions for all query sets. Data points are shown for both Mutated Common Species and Mutated Random Combination query sets.

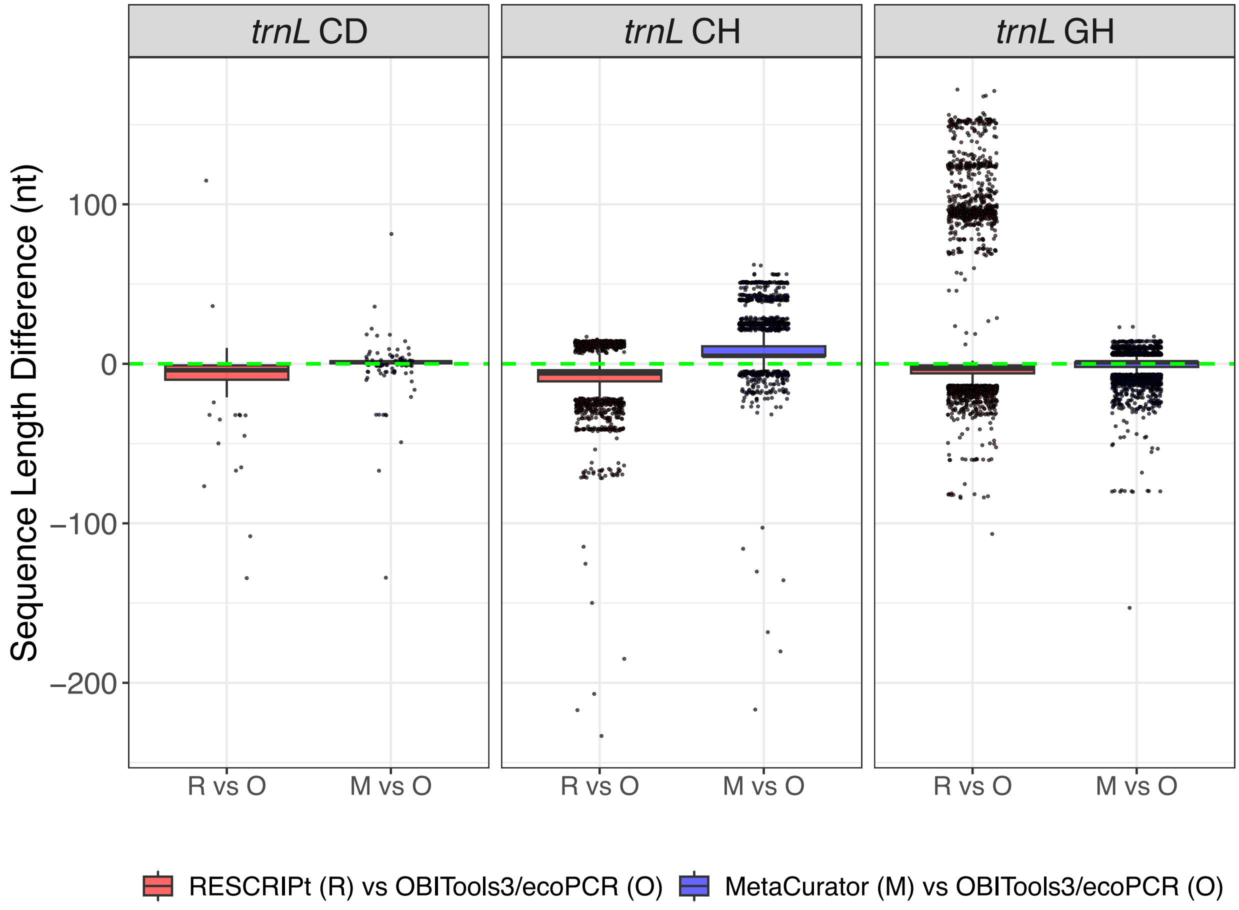


**Supplementary Figure 6. Sequence-length differences for identical accessions across tools for *trnL* CD, CH, and GH regions.**

Sequence-length differences (nucleotides; nt) for identical accessions retrieved by RESCRIPt (**R**) and MetaCurator (**M**) relative to OBITools3/ecoPCR (**O**) for the *trnL* CD, CH, and GH regions. Each panel shows R vs O and M vs O comparisons, with the dashed green line indicating zero difference.

| Input Files | Number of Sequences | Keyword |
|---|---|---|
| **Genbank-1** | 1,058,168 | ("Viridiplantae"[Organism] OR ("Viridiplantae"[Organism] OR Viridiplantae[All Fields])) AND (biomol_genomic[PROP] AND genbank[filter] AND is_nuccore[filter]) AND ("0001/01/01"[PDAT] : "2010/12/31"[PDAT]) |
| **Genbank-2** | 1,612,504 | ("Viridiplantae"[Organism] OR ("Viridiplantae"[Organism] OR Viridiplantae[All Fields])) AND (biomol_genomic[PROP] AND genbank[filter] AND is_nuccore[filter]) AND ("2010/12/30"[PDAT] : "2013/12/31"[PDAT]) |
| **Genbank-3** | 1,717,423 | ("Viridiplantae"[Organism] OR ("Viridiplantae"[Organism] OR Viridiplantae[All Fields])) AND (biomol_genomic[PROP] AND genbank[filter] AND is_nuccore[filter]) AND ("2013/12/30"[PDAT] : "2016/12/31"[PDAT]) |
| **Genbank-4** | 1,130,944 | ("Viridiplantae"[Organism] OR ("Viridiplantae"[Organism] OR Viridiplantae[All Fields])) AND (biomol_genomic[PROP] AND genbank[filter] AND is_nuccore[filter]) AND ("2016/12/30"[PDAT] : "2019/12/31"[PDAT]) |
| **Genbank-5** | 480,821 | ("Viridiplantae"[Organism] OR Viridiplantae[All Fields]) AND (biomol_genomic[PROP] AND genbank[filter] AND is_nuccore[filter] AND ("2019/12/30"[MDAT] : "2020/12/31"[MDAT])) |
| **Genbank-6-a** | 114,318 | ("Viridiplantae"[Organism] OR Viridiplantae[All Fields]) AND (biomol_genomic[PROP] AND genbank[filter] AND is_nuccore[filter] AND ("2020/12/30"[MDAT] : "2021/6/31"[MDAT])) |
| **Genbank-6-b** | 107,850 | ("Viridiplantae"[Organism] OR Viridiplantae[All Fields]) AND (biomol_genomic[PROP] AND genbank[filter] AND is_nuccore[filter] AND ("2021/6/30"[MDAT] : "2021/12/31"[MDAT])) |
| **Genbank-7** | 112,483 | ("Viridiplantae"[Organism] OR Viridiplantae[All Fields]) AND (biomol_genomic[PROP] AND genbank[filter] AND is_nuccore[filter] AND ("2021/12/30"[MDAT] : "2022/6/31"[MDAT])) |
| **Genbank-8-a** | 51,646 | ("Viridiplantae"[Organism] OR Viridiplantae[All Fields]) AND (biomol_genomic[PROP] AND genbank[filter] AND is_nuccore[filter] AND ("2022/6/30"[MDAT] : "2022/9/31"[MDAT])) |
| **Genbank-8-b** | 55,041 | ("Viridiplantae"[Organism] OR Viridiplantae[All Fields]) AND (biomol_genomic[PROP] AND genbank[filter] AND is_nuccore[filter] AND ("2022/9/30"[MDAT] : "2022/12/31"[MDAT])) |
| **Genbank-9** | 153,485 | ("Viridiplantae"[Organism] OR Viridiplantae[All Fields]) AND (biomol_genomic[PROP] AND genbank[filter] AND is_nuccore[filter] AND ("2022/12/30"[MDAT] : "2023/1/31"[MDAT])) |
| **Genbank-10** | 122,498 | ("Viridiplantae"[Organism] OR Viridiplantae[All Fields]) AND (biomol_genomic[PROP] AND genbank[filter] AND is_nuccore[filter] AND ("2023/1/30"[MDAT] : "2023/6/31"[MDAT])) |
| **Genbank-11** | 65,132 | ("Viridiplantae"[Organism] OR Viridiplantae[All Fields]) AND (biomol_genomic[PROP] AND genbank[filter] AND is_nuccore[filter] AND ("2023/6/30"[MDAT] : "2023/9/31"[MDAT])) |
| **Genbank-12** | 78,985 | ("Viridiplantae"[Organism] OR Viridiplantae[All Fields]) AND (biomol_genomic[PROP] AND genbank[filter] AND is_nuccore[filter] AND ("2023/9/30"[MDAT] : "2023/12/31"[MDAT])) |
| **Genbank-13** | 68,674 | ("Viridiplantae"[Organism] OR Viridiplantae[All Fields]) AND (biomol_genomic[PROP] AND genbank[filter] AND is_nuccore[filter] AND ("2023/12/30"[MDAT] : "2024/4/31"[MDAT])) |

| | | |
|---|---|---|
| **Genbank-14** | 37,204 | ("Viridiplantae"[Organism] OR Viridiplantae[All Fields]) AND (biomol_genomic[PROP] AND genbank[filter] AND is_nuccore[filter] AND ("2024/4/30"[MDAT] : "2024/12/31"[MDAT])) |
| **TPA** | 3,014 | ("Viridiplantae"[Organism] OR Viridiplantae[All Fields]) AND (biomol_genomic[PROP] AND tpa_srcdb[filter] AND is_nuccore[filter]) |
| **EMBL** | 4,792,161 | ("Viridiplantae"[Organism] OR Viridiplantae[All Fields]) AND (biomol_genomic[PROP] AND embl[filter] AND is_nuccore[filter]) |
| **DDBJ** | 1,077,244 | ("Viridiplantae"[Organism] OR Viridiplantae[All Fields]) AND (biomol_genomic[PROP] AND ddbj[filter] AND is_nuccore[filter]) |
| **BOLD** | 531,052 | Bryophyta\|Chlorophyta\|Lycopodiophyta\|Magnoliophyta\|Pinophyta\|Pteridophyta |
| **Refseq-1** | 969,137 | ("Viridiplantae"[Organism] OR ("Viridiplantae"[Organism] OR Viridiplantae[All Fields])) AND (biomol_genomic[PROP] AND refseq[filter] AND is_nuccore[filter]) AND ("0001/01/01"[MDAT] : "2018/12/31"[MDAT]) |
| **Refseq-2** | 903,013 | ("Viridiplantae"[Organism] OR Viridiplantae[All Fields]) AND (biomol_genomic[PROP] AND refseq[filter] AND ("2018/12/30"[MDAT] : "2021/12/30"[MDAT])) |
| **Refseq-3** | 198,864 | ("Viridiplantae"[Organism] OR Viridiplantae[All Fields]) AND (biomol_genomic[PROP] AND refseq[filter] AND ("2021/12/29"[MDAT] : "2024/12/30"[MDAT])) |
| **Total** | 15,441,661 | |

**Supplementary Table 1. Keywords used to retrieve nucleotide sequences from INSDC and BOLD, and the number of sequences obtained in each partition.**

| *trnL* Region | Tools | Raw FASTA | Curation Steps<br>Ambiguous Base Removal | Taxonomy Cleanup | Length-based Filtering | Sequence Dereplication |
|---|---|---|---|---|---|---|
| CD | **OBITools3/ecoPCR** | 70,446 | 69,465 | 69,122 | 68,860 | 32,286 |
| | **RESCRIPt** | 1,303,060 | 161,074 | 160,550 | 149,474 | 87,138 |
| | **MetaCurator** | 178,456 | 170,412 | 170,071 | 168,743 | 93,429 |
| CH | **OBITools3/ecoPCR** | 74,017 | 73,573 | 73,444 | 73,279 | 30,351 |
| | **RESCRIPt** | 1,632,729 | 192,998 | 192,570 | 170,071 | 81,995 |
| | **MetaCurator** | 104,719 | 103,074 | 102,852 | 101,721 | 46,844 |
| GH | **OBITools3/ecoPCR** | 225,788 | 224,183 | 223,538 | 222,760 | 88,544 |
| | **RESCRIPt** | 1,568,523 | 144,750 | 144,334 | 144,333 | 68,974 |
| | **MetaCurator** | 133,054 | 131,930 | 131,565 | 128,055 | 59,079 |

**Supplementary Table 2. Number of sequences retained at each curation step.**
Summary of sequence retention across different curation steps for the *trnL* CD, CH, and GH regions using OBITools3/ecoPCR, RESCRIPt, and MetaCurator. The number of entries at each stage represents the sequences remaining after successive filtering steps.

| | *trnL* CD | | | | | *trnL* CH | | | | | *trnL* GH | | | | |
|---|---|---|---|---|---|---|---|---|---|---|---|---|---|---|---|
| **Taxonomic Level** | **Class** | **Order** | **Family** | **Genus** | **Species** | **Class** | **Order** | **Family** | **Genus** | **Species** | **Class** | **Order** | **Family** | **Genus** | **Species** |
| **OBITools3/ecoPCR** | 21 | 122 | 520 | 5263 | 24534 | 22 | 122 | 522 | 5374 | 25460 | 22 | 136 | 665 | 10901 | 75710 |
| **RESCRIPt** | 13 | 102 | 485 | 9183 | 60482 | 16 | 117 | 566 | 9923 | 66136 | 12 | 94 | 440 | 8486 | 57155 |
| **MetaCurator** | 20 | 124 | 580 | 9307 | 63845 | 15 | 123 | 595 | 7474 | 39127 | 14 | 105 | 471 | 8237 | 49812 |
| **Shared between OBITools3/ecoPCR & RESCRIPt** | 13 | 97 | 396 | 4333 | 15771 | 15 | 108 | 464 | 5025 | 22299 | 12 | 93 | 434 | 8308 | 52630 |
| **Shared between OBITools3/ecoPCR & MetaCurator** | 19 | 110 | 456 | 4665 | 21599 | 14 | 110 | 483 | 4793 | 21408 | 14 | 104 | 465 | 8052 | 47168 |
| **Shared between RESCRIPt & MetaCurator** | 12 | 95 | 416 | 8067 | 51645 | 13 | 111 | 516 | 7021 | 36124 | 11 | 89 | 401 | 7187 | 41393 |
| **Shared across all databases** | 12 | 91 | 356 | 3974 | 14462 | 12 | 103 | 439 | 4579 | 19873 | 11 | 88 | 398 | 7108 | 39738 |
| **Unique to OBITools3/ecoPCR** | 1 | 6 | 24 | 239 | 1626 | 5 | 7 | 14 | 135 | 1626 | 7 | 27 | 164 | 1649 | 15650 |
| **Unique to RESCRIPt** | 0 | 1 | 29 | 757 | 7528 | 0 | 1 | 25 | 2456 | 27586 | 0 | 0 | 3 | 99 | 2870 |
| **Unique to MetaCurator** | 1 | 10 | 64 | 549 | 5063 | 0 | 5 | 35 | 239 | 1468 | 0 | 0 | 3 | 106 | 989 |
| **Present in OBITools3/ecoPCR but absent in RESCRIPt** | 8 | 25 | 124 | 930 | 8763 | 7 | 14 | 58 | 349 | 3161 | 10 | 43 | 231 | 2593 | 23080 |
| **Present in OBITools3/ecoPCR but absent in MetaCurator** | 2 | 12 | 64 | 598 | 2935 | 8 | 12 | 39 | 581 | 4052 | 8 | 32 | 200 | 2849 | 28542 |
| **Present in RESCRIPt but absent in MetaCurator** | 1 | 7 | 69 | 1116 | 8837 | 3 | 6 | 50 | 2902 | 30012 | 1 | 5 | 39 | 1299 | 15762 |
| **Present in RESCRIPt but absent in OBITools3/ecoPCR** | 0 | 5 | 89 | 4850 | 44711 | 1 | 9 | 102 | 4898 | 43837 | 0 | 1 | 6 | 178 | 4525 |
| **Present in MetaCurator but absent in OBITools3/ecoPCR** | 1 | 14 | 124 | 4642 | 42246 | 1 | 13 | 112 | 2681 | 17719 | 0 | 1 | 6 | 185 | 2644 |
| **Present in MetaCurator but absent in RESCRIPt** | 8 | 29 | 164 | 1240 | 12200 | 2 | 12 | 79 | 453 | 3003 | 3 | 16 | 70 | 1050 | 8419 |

**Supplementary Table 3. Taxonomic comparison of *trnL* CD, CH and GH reference databases generated by OBITools3/ecoPCR, RESCRIPt, and MetaCurator.**
Number of unique entries at different taxonomic levels (Class, Order, Family, Genus, and Species) for *trnL* CD, CH, and GH regions. The table shows both shared and unique taxonomic entries across tools, including taxa detected by one tool but absent from others.

| | | Taxonomic Levels | | | | | | | | | | | | | | | | | |
|---|---|---|---|---|---|---|---|---|---|---|---|---|---|---|---|---|---|---|---|
| | | Species | | | | | | Genus | | | | | | Family | | | | | |
| **Query Sets** | **DBs** | **CC** | **UC** | **MC** | **FC** | **P** | **R** | **CC** | **UC** | **MC** | **FC** | **P** | **R** | **CC** | **UC** | **MC** | **FC** | **P** | **R** |
| **CS** | **O** | 1,543 | 530 | 927 | 0.823 | 0.625 | 0.514 | 2,567 | 191 | 242 | 0.936 | 0.914 | 0.856 | 2,990 | 3 | 7 | 0.999 | 0.998 | 0.997 |
| **CS** | **R** | 1,660 | 470 | 870 | 0.843 | 0.656 | 0.553 | 2,632 | 158 | 210 | 0.947 | 0.926 | 0.877 | 2,988 | 3 | 9 | 0.999 | 0.997 | 0.996 |
| **CS** | **M** | 1,726 | 432 | 842 | 0.856 | 0.672 | 0.575 | 2,713 | 105 | 182 | 0.965 | 0.937 | 0.904 | 2,991 | 0 | 9 | 1.000 | 0.997 | 0.997 |
| **MCS** | **O** | 1,461 | 850 | 689 | 0.717 | 0.680 | 0.487 | 2,587 | 200 | 213 | 0.933 | 0.924 | 0.862 | 2,990 | 3 | 7 | 0.999 | 0.998 | 0.997 |
| **MCS** | **R** | 1,443 | 937 | 620 | 0.688 | 0.699 | 0.481 | 2,600 | 233 | 167 | 0.922 | 0.940 | 0.867 | 2,988 | 3 | 9 | 0.999 | 0.997 | 0.996 |
| **MCS** | **M** | 1,442 | 933 | 625 | 0.689 | 0.698 | 0.481 | 2,639 | 201 | 160 | 0.933 | 0.943 | 0.880 | 2,983 | 7 | 10 | 0.998 | 0.997 | 0.994 |
| **RC** | **O** | 1,242 | 651 | 1,107 | 0.783 | 0.529 | 0.414 | 2,521 | 209 | 270 | 0.930 | 0.903 | 0.840 | 2,993 | 3 | 4 | 0.999 | 0.999 | 0.998 |
| **RC** | **R** | 1,439 | 564 | 997 | 0.812 | 0.591 | 0.480 | 2,624 | 157 | 219 | 0.948 | 0.923 | 0.875 | 2,988 | 3 | 9 | 0.999 | 0.997 | 0.996 |
| **RC** | **M** | 1,506 | 520 | 974 | 0.827 | 0.607 | 0.502 | 2,636 | 139 | 225 | 0.954 | 0.921 | 0.879 | 2,985 | 9 | 6 | 0.997 | 0.998 | 0.995 |
| **MRC** | **O** | 1,161 | 963 | 876 | 0.679 | 0.570 | 0.387 | 2,561 | 230 | 209 | 0.923 | 0.925 | 0.854 | 2,993 | 4 | 3 | 0.999 | 0.999 | 0.998 |
| **MRC** | **R** | 1,271 | 1,020 | 709 | 0.660 | 0.642 | 0.424 | 2,624 | 226 | 150 | 0.925 | 0.946 | 0.875 | 2,989 | 3 | 8 | 0.999 | 0.997 | 0.996 |
| **MRC** | **M** | 1,334 | 967 | 699 | 0.678 | 0.656 | 0.445 | 2,596 | 240 | 164 | 0.920 | 0.941 | 0.865 | 2,979 | 12 | 9 | 0.996 | 0.997 | 0.993 |

**Supplementary Table 4. Taxonomic classification metrics across all query sets and databases for the *trnL* CD region.**
Comparison of taxonomic classification performance at the species, genus, and family levels across different query sets and databases. Metrics include the number of correctly classified (**CC**), unclassified (**UC**), and misclassified (**MC**) entries, along with fraction classified (**FC**), precision (**P**), and recall (**R**). Query sets: Common Species (**CS**), Mutated Common Species (**MCS**), Random Combination (**RC**), Mutated Random Combination (**MRC**). Databases (**DBs**): OBITools3/ecoPCR (**O**), RESCRIPt (**R**), MetaCurator (**M**).

| | | Taxonomic Levels | | | | | | | | | | | | | | | | | |
|---|---|---|---|---|---|---|---|---|---|---|---|---|---|---|---|---|---|---|---|
| | | Species | | | | | | Genus | | | | | | Family | | | | | |
| Query Sets | DBs | CC | UC | MC | FC | P | R | CC | UC | MC | FC | P | R | CC | UC | MC | FC | P | R |
| CS | O | 734 | 1,613 | 653 | 0.462 | 0.529 | 0.245 | 1,655 | 1,100 | 245 | 0.633 | 0.871 | 0.552 | 2,924 | 67 | 9 | 0.978 | 0.997 | 0.975 |
| CS | R | 690 | 1,621 | 689 | 0.460 | 0.500 | 0.230 | 1,620 | 1,170 | 210 | 0.610 | 0.885 | 0.540 | 2,875 | 106 | 19 | 0.965 | 0.993 | 0.958 |
| CS | M | 652 | 2,050 | 298 | 0.317 | 0.686 | 0.217 | 1,548 | 1,329 | 123 | 0.557 | 0.926 | 0.516 | 2,970 | 20 | 10 | 0.993 | 0.997 | 0.990 |
| MCS | O | 524 | 2,119 | 357 | 0.294 | 0.595 | 0.175 | 1,429 | 1,385 | 186 | 0.538 | 0.885 | 0.476 | 2,884 | 108 | 8 | 0.964 | 0.997 | 0.961 |
| MCS | R | 499 | 2,071 | 430 | 0.310 | 0.537 | 0.166 | 1,384 | 1,433 | 183 | 0.522 | 0.883 | 0.461 | 2,830 | 161 | 9 | 0.946 | 0.997 | 0.943 |
| MCS | M | 455 | 2,352 | 193 | 0.216 | 0.702 | 0.152 | 1,260 | 1,626 | 114 | 0.458 | 0.917 | 0.420 | 2,902 | 90 | 8 | 0.970 | 0.997 | 0.967 |
| RC | O | 628 | 1,640 | 732 | 0.453 | 0.462 | 0.209 | 1,712 | 1,077 | 211 | 0.641 | 0.890 | 0.571 | 2,910 | 76 | 14 | 0.975 | 0.995 | 0.970 |
| RC | R | 637 | 1,618 | 745 | 0.461 | 0.461 | 0.212 | 1,721 | 1,063 | 216 | 0.646 | 0.888 | 0.574 | 2,899 | 84 | 17 | 0.972 | 0.994 | 0.966 |
| RC | M | 507 | 2,209 | 284 | 0.264 | 0.641 | 0.169 | 1,320 | 1,541 | 139 | 0.486 | 0.905 | 0.440 | 2,711 | 268 | 21 | 0.911 | 0.992 | 0.904 |
| MRC | O | 484 | 2,075 | 441 | 0.308 | 0.523 | 0.161 | 1,479 | 1,347 | 174 | 0.551 | 0.895 | 0.493 | 2,860 | 129 | 11 | 0.957 | 0.996 | 0.953 |
| MRC | R | 473 | 2,090 | 437 | 0.303 | 0.520 | 0.158 | 1,514 | 1,313 | 173 | 0.562 | 0.897 | 0.505 | 2,854 | 133 | 13 | 0.956 | 0.995 | 0.951 |
| MRC | M | 361 | 2,453 | 186 | 0.182 | 0.660 | 0.120 | 1,062 | 1,808 | 130 | 0.397 | 0.891 | 0.354 | 2,648 | 334 | 18 | 0.889 | 0.993 | 0.883 |

**Supplementary Table 5. Taxonomic classification metrics across all query sets and databases for the *trnL* CH region.**
Comparison of taxonomic classification performance at the species, genus, and family levels across different query sets and databases. Metrics include the number of correctly classified (**CC**), unclassified (**UC**), and misclassified (**MC**) entries, along with fraction classified (**FC**), precision (**P**), and recall (**R**). Query sets: Common Species (**CS**), Mutated Common Species (**MCS**), Random Combination (**RC**), Mutated Random Combination (**MRC**). Databases (**DBs**): OBITools3/ecoPCR (**O**), RESCRIPt (**R**), MetaCurator (**M**).

| | | Taxonomic Levels | | | | | | | | | | | | | | | | | |
|---|---|---|---|---|---|---|---|---|---|---|---|---|---|---|---|---|---|---|---|
| | | Species | | | | | | Genus | | | | | | Family | | | | | |
| **Query Sets** | **DBs** | **CC** | **UC** | **MC** | **FC** | **P** | **R** | **CC** | **UC** | **MC** | **FC** | **P** | **R** | **CC** | **UC** | **MC** | **FC** | **P** | **R** |
| **CS** | **O** | 234 | 2,539 | 227 | 0.154 | 0.508 | 0.078 | 662 | 2,221 | 117 | 0.260 | 0.850 | 0.221 | 1,470 | 1,517 | 13 | 0.494 | 0.991 | 0.490 |
| **CS** | **R** | 192 | 2,528 | 280 | 0.157 | 0.407 | 0.064 | 718 | 2,161 | 121 | 0.280 | 0.856 | 0.239 | 1,587 | 1,406 | 7 | 0.531 | 0.996 | 0.529 |
| **CS** | **M** | 364 | 2,385 | 251 | 0.205 | 0.592 | 0.121 | 901 | 1,984 | 115 | 0.339 | 0.887 | 0.300 | 1,808 | 1,185 | 7 | 0.605 | 0.996 | 0.603 |
| **MCS** | **O** | 184 | 2,638 | 178 | 0.121 | 0.508 | 0.061 | 584 | 2,292 | 124 | 0.236 | 0.825 | 0.195 | 1,479 | 1,512 | 9 | 0.496 | 0.994 | 0.493 |
| **MCS** | **R** | 153 | 2,607 | 240 | 0.131 | 0.389 | 0.051 | 660 | 2,208 | 132 | 0.264 | 0.833 | 0.220 | 1,564 | 1,429 | 7 | 0.524 | 0.996 | 0.521 |
| **MCS** | **M** | 320 | 2,490 | 190 | 0.170 | 0.627 | 0.107 | 793 | 2,114 | 93 | 0.295 | 0.895 | 0.264 | 1,781 | 1,212 | 7 | 0.596 | 0.996 | 0.594 |
| **RC** | **O** | 227 | 2,502 | 271 | 0.166 | 0.456 | 0.076 | 832 | 2,068 | 100 | 0.311 | 0.893 | 0.277 | 1,506 | 1,481 | 13 | 0.506 | 0.991 | 0.502 |
| **RC** | **R** | 177 | 2,500 | 323 | 0.167 | 0.354 | 0.059 | 852 | 2,069 | 79 | 0.310 | 0.915 | 0.284 | 1,598 | 1,394 | 8 | 0.535 | 0.995 | 0.533 |
| **RC** | **M** | 301 | 2,433 | 266 | 0.189 | 0.531 | 0.100 | 969 | 1,936 | 95 | 0.355 | 0.911 | 0.323 | 1,782 | 1,210 | 8 | 0.597 | 0.996 | 0.594 |
| **MRC** | **O** | 185 | 2,588 | 227 | 0.137 | 0.449 | 0.062 | 765 | 2,130 | 105 | 0.290 | 0.879 | 0.255 | 1,509 | 1,480 | 11 | 0.507 | 0.993 | 0.503 |
| **MRC** | **R** | 140 | 2,596 | 264 | 0.135 | 0.347 | 0.047 | 788 | 2,123 | 89 | 0.292 | 0.899 | 0.263 | 1,580 | 1,411 | 9 | 0.530 | 0.994 | 0.527 |
| **MRC** | **M** | 275 | 2,529 | 196 | 0.157 | 0.584 | 0.092 | 893 | 2,027 | 80 | 0.324 | 0.918 | 0.298 | 1,742 | 1,249 | 9 | 0.584 | 0.995 | 0.581 |

**Supplementary Table 6. Taxonomic classification metrics across all query sets and databases for the *trnL* GH region.**
Comparison of taxonomic classification performance at the species, genus, and family levels across different query sets and databases. Metrics include the number of correctly classified (**CC**), unclassified (**UC**), and misclassified (**MC**) entries, along with fraction classified (**FC**), precision (**P**), and recall (**R**). Query sets: Common Species (**CS**), Mutated Common Species (**MCS**), Random Combination (**RC**), Mutated Random Combination (**MRC**). Databases (**DBs**): OBITools3/ecoPCR (**O**), RESCRIPt (**R**), MetaCurator (**M**).
